# Lipid-associated architecture and divergent ammonium transport distinguish a bacterial Rhesus protein

**DOI:** 10.64898/2026.09.17.752382

**Authors:** Benjamin F. Cooper, Adriana Bizior, Reza Talandashti, Abraham O. Oluwole, Peter Henderson, Thomas Harris, Carol V. Robinson, Paul A. Hoskisson, Syma Khalid, Leighton Pritchard, Georgia L. Isom, Arnaud Javelle

**Author notes:** These authors contributed equally to this work (Benjamin F. Cooper, Adriana Bizior).

## Abstract

Nitrification is a major process in the global nitrogen cycle, initiated by the oxidation of ammonia to nitrite. The ammonia-oxidising bacterium *Nitrosomonas europaea* catalyses this first and rate-limiting step and depends on ammonium both as an energy substrate and as a nitrogen source, making ammonium acquisition central to its physiology. Cellular ammonium transport is mediated by the Amt/Mep/Rh superfamily, whose members share a conserved fold despite divergent physiological roles. Remarkably, *N. europaea* lacks a canonical prokaryotic Amt transporter and instead expresses NeRh50, a Rh-family protein proposed to have been acquired from a eukaryotic lineage. Here, we determine a cryo-EM structure of NeRh50 that reveals tightly associated lipids at the monomer–monomer interfaces, including density consistent with an unusual inverted lipid orientation. Native mass spectrometry and molecular dynamics simulations further support persistent lipid association at these interfaces. NeRh50 also differs functionally from canonical AmtB, selectively mediating electrogenic ammonium transport with ∼70-fold lower apparent affinity. Transport is enhanced under alkaline conditions but not by an imposed proton gradient, suggesting membrane-potential dependence. Together, these findings reveal divergence from canonical Amt proteins in both membrane-associated architecture and transport properties, providing insight into the functional and evolutionary diversification of the Amt/Mep/Rh superfamily.

## Introduction

The exchange of ammonium across cellular membranes is a fundamental process in all domains of life. For bacteria, fungi and plants, ammonium is a preferred nitrogen source acquired from the environment, whereas for mammals it represents a cytotoxic metabolic waste product that must be efficiently excreted^1^. This physiological duality is served by the ubiquitous Amt/Mep/Rh superfamily, comprising prokaryotic ammonium transporters (Amt), fungal methylammonium permeases (Mep) and Rhesus (Rh) proteins^2^. Rh proteins are abundant in vertebrates, where they are expressed in erythrocytes, the kidneys and the liver^3^, but also occur in ammonia-oxidising bacteria and a small number of archaea^4^. Despite their markedly different physiological roles, Amt/Mep/Rh proteins share a highly conserved fold. Structures of various Amt, Mep and Rh across the superfamily reveal a common trimeric architecture in which each monomer comprises 11 transmembrane helices^5^. The structural basis of this functional diversification is therefore unlikely to be explained by the conserved scaffold alone and requires consideration of transport behaviour, conformational dynamics and interactions with the surrounding membrane^6^.

A decisive advance in understanding Amt transport was provided by solid-supported membrane electrophysiology (SSME), which established electrogenic ammonium transport, first for *Archaeoglobus fulgidus* Amts and subsequently for *E. coli* AmtB^7,8^. Based on these data, Williamson *et al.* proposed a two-lane mechanism in which NH_4_^+^ is deprotonated at the periplasmic face, with NH_3_ and H^+^ traversing spatially distinct routes through the pore^8^. More recently, kinetic and thermodynamic modelling supported the requirement for membrane-potential-driven transport to achieve active ammonium accumulation and described a spatiotemporal mechanism coupling ammonia and proton fluxes through AmtB^9^. Both frameworks therefore place electrogenicity and membrane potential at the centre of Amt transport, although the precise molecular mechanism remains under investigation^4,6^.

Among prokaryotic Rh proteins, NeRh50 from *Nitrosomonas europaea* provides a particularly informative system in which to examine how the Rh branch has diverged from canonical Amt transporters. *N. europaea* is an obligate ammonia-oxidising bacterium that catalyses the first and rate-limiting step of nitrification and depends on ammonium both as an energy substrate and as a nitrogen source. Remarkably, its genome encodes no canonical Amt transporter; instead, its only identified ammonium transporter is NeRh50, a Rh-family protein proposed to have been acquired from a eukaryotic lineage^10^. Crystal structures established that NeRh50 retains the conserved Rh fold while revealing differences from AmtB, including the absence of the canonical periplasmic Am1/S1 ammonium-binding site and a more open Phe-gate associated with a wider pore entrance^11^. However, the intrinsic transport properties of purified wild-type NeRh50 remain incompletely defined, leaving unresolved how these structural differences translate into Rh-family transport.

The membrane environment provides a second potential dimension of divergence. Specific lipid interactions modulate AmtB conformation, dynamics and electrogenic transport, with phospholipids occupying sites that include the monomer–monomer interface^12,13,14^. Whether comparable interfacial lipid interactions occur in Rh-family transporters remains unknown, and no associated phospholipid density has been resolved in previous NeRh50 structures^11^. Thus, both the membrane-associated architecture and intrinsic transport properties of NeRh50 remain poorly defined.

Here, we combine cryo-electron microscopy, native mass spectrometry and molecular dynamics simulations to investigate the lipid-associated architecture of NeRh50, and use SSME to define its substrate selectivity, apparent affinity, pH dependence and proton-gradient dependence. Together, our findings reveal that a deeply conserved membrane-protein scaffold can accommodate striking divergence in both its interactions with the surrounding membrane and its transport properties, establishing membrane-associated architecture and transport behaviour as complementary dimensions of functional diversification within the Amt/Mep/Rh superfamily.

## Results

### Cryo-EM structure of NeRh50

To investigate the architecture of NeRh50 free from crystal packing constraints we determined its structure via cryo-EM at a global resolution of 3.1 Å (Figure 1A, Figure S1, Figure S4, Supplementary Table 1). The density was well-resolved throughout the transmembrane region, allowing complete characterisation of the protein’s trimeric core, however, the C-terminal, helical moiety displayed substantial conformational heterogeneity limiting our ability to model the final portion of the protein (residues 410 - 425) (Figure 1A-B). Nevertheless, our cryo-EM model is in agreement with previously published crystallographic NeRh50 structures^11,15^, comprising the canonical trimeric 11-transmembrane-helix fold of the Amt/Mep/Rh superfamily and C-terminal coiled coil, whilst lacking the aromatic-rich Am1/S1 ammonium-binding site of *E. coli* AmtB. Masked refinement focussing upon the core transmembrane region yielded only a marginal improvement in resolution, from 3.1 to 3 Å, whilst the additional application of C3 symmetry resulted in a similarly modest increase to 2.9 Å. Nonetheless, the C3 reconstruction exhibited reduced map quality and connectivity within several loop regions, indicating local deviations from strict threefold symmetry and supporting a pseudosymmetrical arrangement of the complex (Figure S1, Supplementary Table 1).

**Figure 1.**
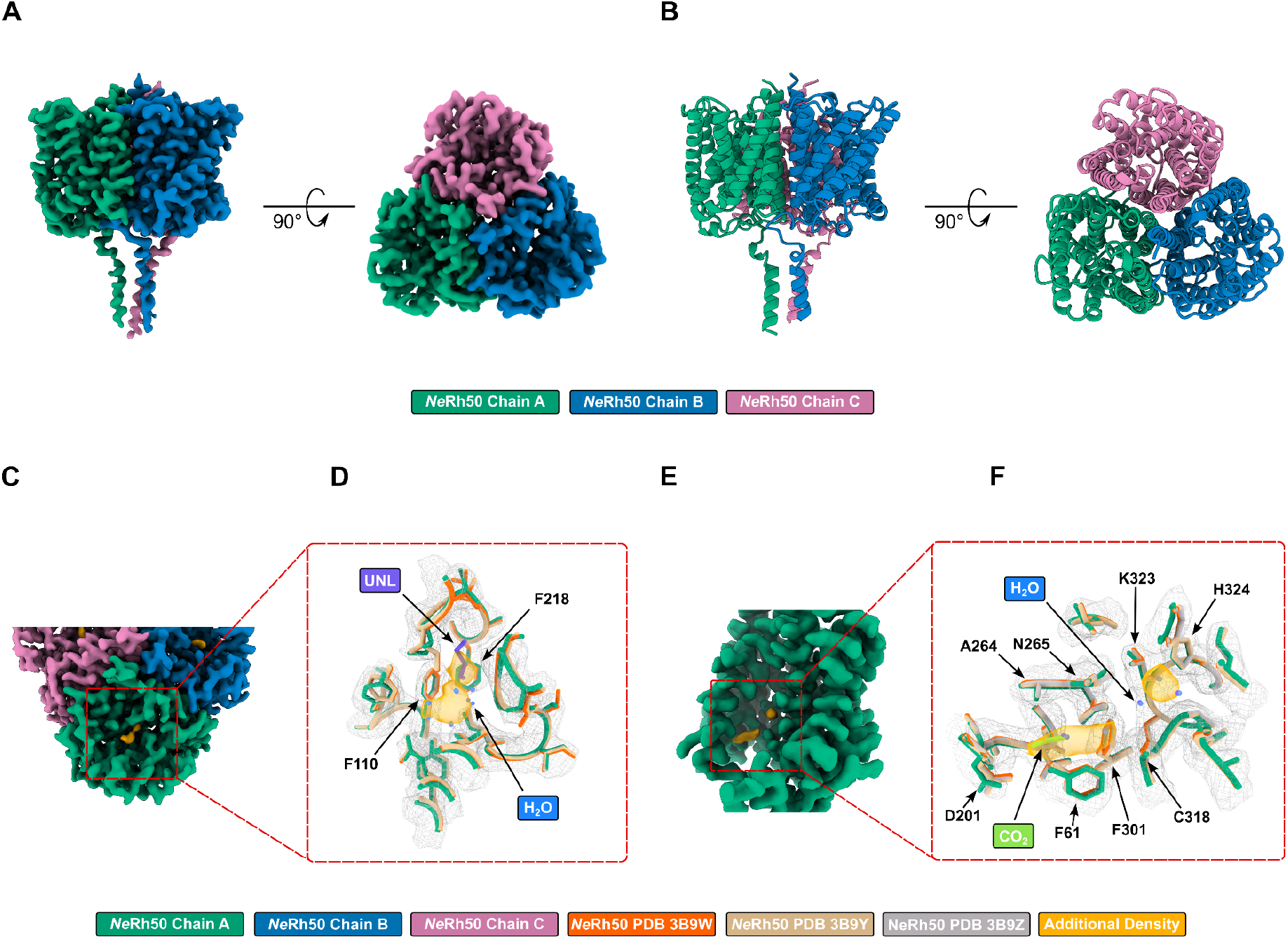
Cryo-EM Reconstruction of NeRh50. **(A)** LAFTER^16^ filtered NeRh50 cryo-EM reconstruction at 3.1 A. Density corresponding to chains A, B and C are coloured green, blue and pink respectively. **(B)** Atomic model built from the NeRh50 cryo-EM reconstruction. Chains A, B and C are coloured green, blue and pink respectively. **(C)** Elongated density (gold) present within the periplasmic vestibule of NeRh50. The Relion postprocessed map is contoured at 1.5 σ (map level 0.008), with density corresponding to chains A, B and C coloured green, blue and pink respectively. **(D)** Density (gold) present within the periplasmic vestibule of NeRh50. The Relion postprocessed map contoured at 1.5 σ (map level 0.008) and displayed as a grey mesh. NeRh50 models derived from our cryo-EM data, PDB 3B9W and PDB 3B9Y are coloured green, orange and tan, respectively. Water molecules modelled in both PDBs 3B9W and 3B9Y are coloured blue, whilst the unknown ligand modelled in PDB 3B9Y is coloured purple. **(E)** Elongated density (gold) present within the cytoplasmic vestibule of NeRh50. The Relion postprocessed map is contoured at 1.5 σ (map level 0.008), with density corresponding to chain A coloured green. Density corresponding to residues 32-82 (TM1-2) of NeRh50 is removed to aid clarity. **(F)** Densities (gold) present within the cytoplasmic vestibule of NeRh50. The Relion postprocessed map is contoured at 1.5 σ (map level 0.008) and displayed as a grey mesh. NeRh50 models derived from our cryo-EM data, PDB 3B9W, PDB 3B9Y and PDB 3B9Z are coloured green, orange, tan and grey, respectively. Water molecules modelled in both PDBs 3B9W and 3B9Y are coloured blue, whilst the CO_2_ molecule modelled in PDB 3B9Z is coloured lime green.

All three maps contained additional densities not accounted for by the modelled protein moieties, several of which were located within or adjacent to the central pore of each monomer. Elongated densities were observed within the periplasmic vestibule of each protomer, apparently blocked from entering the central pore by the highly conserved, gating residue Phe218 (Figure 1C-D, Figure S2). Analogous densities are present in the existing crystallographic structures of NeRh50, being modelled as a series of water molecules and an unknown ligand in PDB entries 3B9W and 3B9Y, respectively (Figure 1D, Figure S2). The recurrence of these densities across cryo-EM and crystal structures suggests the presence of a tightly associated ligand or network of water molecules, however, their identity cannot be unambiguously assigned at the resolution of our map.

Two additional densities were observed towards the cytoplasmic vestibule of each protomer. A smaller globular density was positioned at the cytoplasmic entrance to the pore, approximately equidistant between His324 and Cys318 (Figure 1E, Figure S2). His324 forms the lower component of the central twin-His motif, whereas Cys318, together with Lys323, marks the end of the NeRh50 hydrophobic pore. Once again, this density aligns well with a series of water molecules modelled in PDB entries 3B9W and 3B9Y (Figure 1E, Figure S2). A second elongated density was observed slightly lower in the cytoplasmic vestibule, adjacent to the pore mouth. Intriguingly, this density partially overlaps with a putative CO_2_ binding pocket, previously identified in a crystallographic structure of CO_2_-pressurised NeRh50 (PDB 3B9Z) (Figure 1E, F, Figure S2). Nevertheless, the density in our cryo-EM map extends beyond this region towards Cys318 and the pore entry, overlapping water molecules modelled in PDB entries 3B9W and 3B9Y (Figure 1E, F, Figure S2).

### NeRh50 retains tightly bound annular lipids, particularly at the monomer-monomer interface

In addition to the vestibules, non-protein densities were observed around the periphery of the protein, most likely resulting from annular lipid or detergent molecules (Figure 2A). These densities were best resolved at the interface between NeRh50 monomers, particularly within the cleft formed by the packing of the M2 and M4 helices of one monomer against the M6 and M7 helices of the adjacent monomer (Figure 2B, Figure S3). This cleft is occupied by two β-OG detergent molecules in the existing crystallographic structure of NeRh50 (Figure 2C, Figure S3) suggesting this site might harbour amphipathic molecules more stably than the rest of the protein’s exterior.

**Figure 2.**
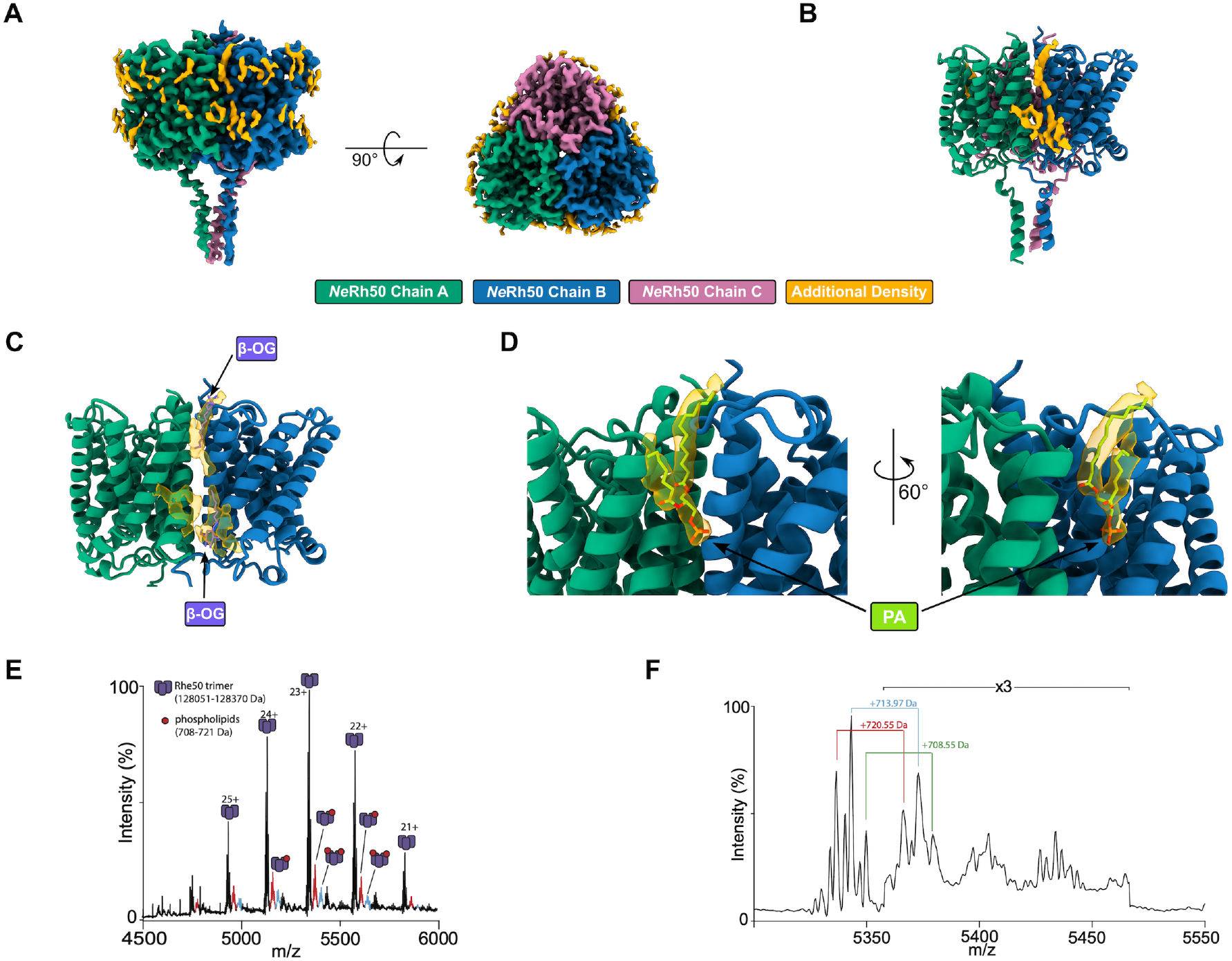
Annular densities surround the NeRh50 transmembrane region. **(A)** Annular densities (gold) surrounding the NeRh50 transmembrane region. The Relion postprocessed map is contoured at 1 σ (map level 0.005), with density corresponding to chains A, B and C coloured green, blue and pink respectively. **(B)** Annular densities (gold) located at the NeRh50 monomer interface between chains A and B (green and blue cartoons respectively). **(C)** Interface density (transparent gold) between NeRh50 chains A and B (green and blue cartoons, respectively) overlaid with the two β-OG molecules (purple) modelled in the crystallographic structure of NeRh50 (PDB 3B9Y). **(D)** Phosphatidic acid (PA, lime green) modelled into the periplasmic leaflet interface density (transparent gold) between NeRh50 chains A and B (green and blue cartoons, respectively). **(E)** Native mass spectrum of NeRh50. Main charge states correspond to NeRh50 trimer in apo- and phospholipid-bound forms. Protein ions were released into the mass spectrometer from β-OG micelles using an activation voltage 100 V. **(F)** Zoomed view of a charge state showing the heterogeneity of the apo protein peaks, spaced by ∼158 Da, in addition to 64-Da Ni2+ adducts.

In all three of our NeRh50 cryo-EM maps, the additional density towards the periplasmic side of the cleft was particularly well defined and exhibited a shape more reminiscent of a glycerophospholipid rather than either of the detergents utilised during the purification protocol (LDAO and DDM). Nevertheless, we were unable to model either of the major *E. coli* lipid species, phosphatidylethanolamine or phosphatidylglycerol, due to the absence of density for the head group moieties of these lipids. Only phosphatidic acid, the universal lipid precursor, was able to reasonably fit the density owing to its small phosphomonoester head group (Figure 2D, Figure S3). Intriguingly, if this density was to arise from a tightly coordinated lipid species, then its orientation would be inverted within the membrane with no obvious way of stabilising its head group within the hydrophobic membrane core. Given our inability to confidently assign individual lipid species from the cryo-EM maps alone, we used native mass spectrometry to investigate whether phospholipids co-purify with NeRh50. The resulting spectra exhibited charge states corresponding to the NeRh50 trimer, together with additional adduct peaks at mass increments of ∼700 Da, consistent with bound phospholipids (Figure 2E). The apo peak also exhibited fine structure attributable to the loss of one to two terminal amino acid residues and adduction with divalent cations, most likely Co^2+^ from the affinity purification (Figure 2F). Accounting for the heterogeneity of the observed masses, the lipid adducts are most consistent with phosphatidylethanolamine (PE) and/or phosphatidylglycerol (PG) species.

To further probe the significance of the lipid species captured by cryo-EM, we used molecular dynamics (MD) to simulate the transmembrane region of NeRh50 embedded in a lipid bilayer that mimicked the composition of the *E. coli* inner membrane (75% POPE, 20% POPG, and 5% cardiolipin) from which it was purified (Figure 3A). Lipid–protein interactions were analysed from the MD trajectories using PyLipID^17^, with particular emphasis on lipid residence time as a measure of the kinetic stability of lipid–protein interactions. Long residence times were predominantly associated with residues located at the monomer interfaces, with several lipid interactions persisting for up to the full 1 µs duration (Figure 3B-C). Within the conditions and timeframe tested, this interfacial binding did not appear to be specific to a single lipid species, though POPE was observed most often due its higher abundance.

**Figure 3.**
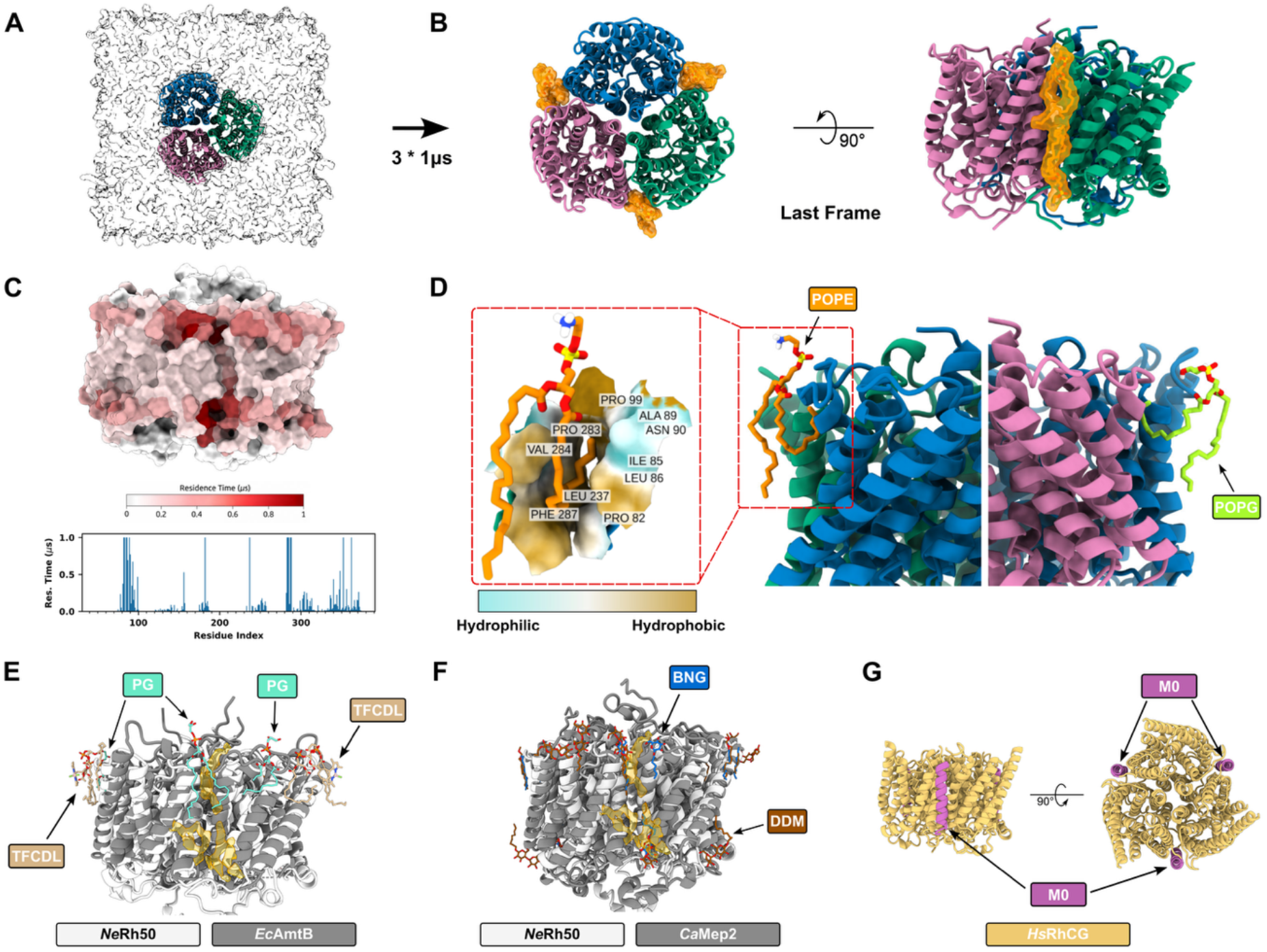
Molecular dynamics simulations of the NeRh50 transmembrane region in a model Gram-negative bacterial inner membrane. **(A)** Overview of the molecular dynamics (MD) simulation setup. The transmembrane region of NeRh50 was embedded in a model Gram-negative bacterial inner membrane composed of POPE, POPG, and cardiolipin at a 75:20:5 ratio. The protein is shown in cartoon representation, with chains A, B and C coloured green, blue and pink respectively, and the membrane is shown as a transparent surface. Three independent 1-µs simulations were performed, yielding a total simulation time of 3 µs. **(B)** Top and side views of the protein and associated lipids in the final frame of replica 1. Protein-associated lipids are shown as orange sticks with a transparent surface representation. **(C)** Lipid residence times mapped onto the protein surface. Residence times were averaged over the three independent simulations. The protein surface is coloured from white to red according to increasing lipid residence time. The residue-wise profile (bottom) shows the corresponding lipid residence time as a function of residue index, highlighting regions with prolonged lipid–protein interactions. **(D)** Representative binding poses of POPE and POPG at the interfaces between NeRh50 chains. POPE and POPG are shown as orange and green sticks, respectively, and the protein chains are shown in cartoon representation. The enlarged view shows a representative POPE-binding pose, with the pocket surface coloured according to residue hydrophobicity, ranging from hydrophilic (cyan) to hydrophobic (brown). In addition to these hydrophobic contacts, ASN90 forms a polar interaction with the glycerol backbone of POPE. **(E)** AmtB lipid binding sites observed in crystallographic studies. TopFluor-cardiolipin (TFCDL) and phosphatidylglycerol (PG) are depicted in tan and teal respectively. (PDBs 4NH2 and 6B21). **(F)** Mep2 lipid binding sites observed in crystallographic studies. DDM and nonyl-D-glucopyranoside (BNG) are depicted in brown and blue respectively. (PDBs 5AEZ and 5AH3). **(G)** Cartoon depiction of HsRhCG (PDB 3HD6) highlighting the position of the M0 helix (lilac), which masks the analogous monomer interface site.

A recurrent feature of the observed binding mode was the bending of one lipid acyl chain and its insertion into a pocket between the monomers (Figure 3D), likely due to the predominantly hydrophobic character of the pocket forming several hydrophobic contacts with the acyl chain. In addition, Asn90 forms a polar interaction with the glycerol backbone of POPE. Notably, the orientation of the lipid observed in the cryo-EM structure was not recapitulated in the MD simulations. This difference may reflect the ability of the cryo-EM structure to capture a lipid conformation that is not favoured in a membrane bilayer and thus would not occur spontaneously during MD simulations. Alternatively, the inverted lipids could be an artefact of detergent extraction during cryo-EM sample preparation. Nevertheless, both approaches provide independent evidence that the monomer interface of NeRh50 can stably engage phospholipids.

We considered the possibility that coordination of amphipathic molecules at the monomer interface was a general feature of the AmtB/Mep/Rh superfamily. Existing crystal structures of *E. coli* AmtB in complex with phosphatidylglycerol (PDB 4NH2) and TopFluor-cardiolipin (PDB 6B21) provided some precedent for interfacial lipid binding. TopFluor-cardiolipin was coordinated at distinct sites towards the apex of each AmtB monomer, whereas phosphatidylglycerol occupied multiple sites, including one directly at the monomer– monomer interface (Figure 3E)^12,18^. Whilst no structures of Mep family proteins with bound lipids were available, those of *C. albicans* MEP2 (PDBs 5AEZ and 5FUF) revealed numerous detergent molecules around the protein’s periphery, several of which occupied the monomer-interface cleft (Figure 3F)^19^. Thus, the binding of amphipathic molecules at the monomer interface may represent a widespread characteristic of Amt/Mep/Rh proteins, potentially contributing to their stability and/or function. Finally, *H. sapiens* RhCG comprises an additional TM helix at its N-terminus (M0) which lies within the monomer interface, abolishing the interface cleft completely (Figure 3G)^20^, thus the M0 helix may represent an alternative method of stabilising the trimeric Rh architecture.

Collectively, our cryo-EM, native mass spectrometry data, and MD simulations indicate that NeRh50 forms intimate interactions with membrane phospholipids that persist throughout protein purification. We propose the most tightly associated lipids to reside at the NeRh50 monomer interface, potentially facilitating protein function and/or stability.

### NeRh50 is an electrogenic ammonium transporter

The architectural and lipid-associated distinctions described above when comparing AmtB and NeRh50 may have important implications for the transport mechanism. Most notably, the reshaped extracellular vestibule, altered pore entrance, and potential inverted interfacial lipid at the monomer–monomer junction suggest that NeRh50 differs from canonical Amt transporters at the point where substrate recognition and entry into the conduction pathway are organised^21^. Importantly, this interfacial lipid occupies a region previously identified as a critical determinant of AmtB transport activity^13,22^. These observations provide the rationale for the SSME characterisation that follows, which directly defines the transport activity, electrogenicity, and energetic coupling of NeRh50.

A 200 mM ammonium pulse applied to *E. coli* polar lipid/POPC 2:1 (w/w) proteoliposomes containing NeRh50 reconstituted at a lipid to protein ratio (LPR) of 5 elicited a transient current of approximately 2 nA that decayed back to baseline. No current was detected in protein-free liposomes, confirming that the signal was protein-specific (Figure 4A). To determine whether this signal reflected a complete translocation cycle rather than a substrate-binding event, the maximum amplitude and decay rate constant (k) was measured at three different LPRs. The maximum amplitude and k value increased with protein density in the membrane (LPR 5, 10 and 50, all pairwise comparisons Šidák-corrected p < 0.001), confirming LPR-dependent decay and therefore electrogenic substrate translocation^23^ (Figure 4B-C). Some of these SSME measurements, also reported by Williamson *et al.*^8^, were repeated here so that all subsequent functional measurements could be compared within consistent assay conditions throughout. The current amplitude observed here was in the same low-nA range as reported by Williamson *et al.* The fitted decay constants differed modestly: Williamson *et al.* reported k values of 39.0 ± 3.6 s⁻¹ at LPR 5 and 24.0 ± 1.7 s⁻¹ at LPR 10. Small differences between independent SSME series are not unexpected, as total current can vary between sensors even when the underlying transport behaviour remains highly consistent^24^. Importantly, in both datasets, WT NeRh50 displays clear LPR-dependent current decay, consistent with electrogenic transport rather than a substrate-binding event.

**Figure 4.**
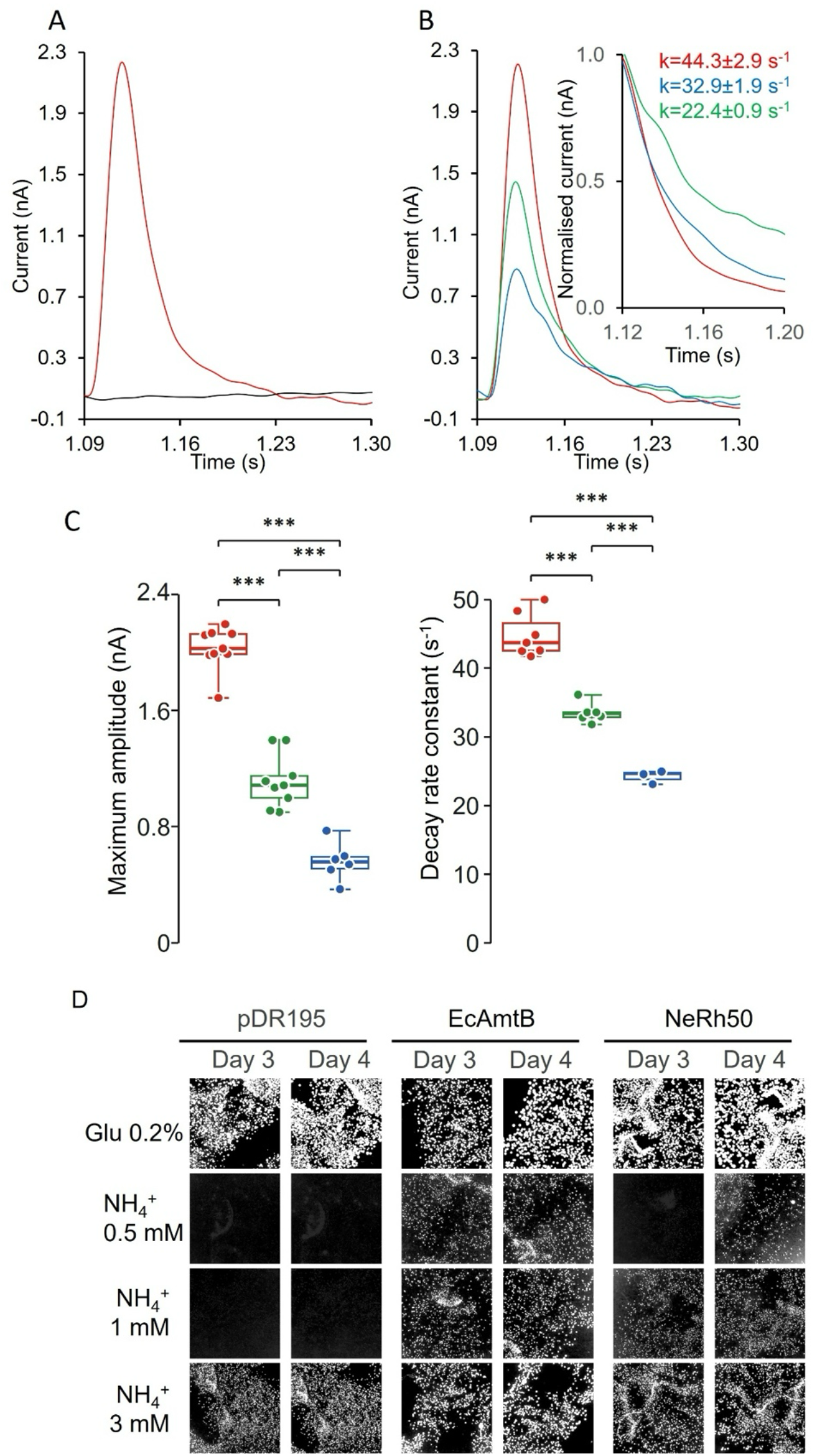
NeRh50 is an electrogenic ammonium transporter. **(A)** Representative SSME transient current recorded after application of NH₄⁺ to proteoliposomes containing NeRh50 at LPR5 (red), with protein-free liposomes shown as a control (black). **(B)** Transient currents recorded at different lipid-to-protein ratios (LPRs) 5 (red), 10 (green) and 50 (blue), with corresponding normalised traces shown in the inset. **(C)** Maximum amplitude and decay rate constant k at each LPR. Individual data points are overlaid on box plots (median, interquartile range, range); LPR5 in red, LPR10 in green, LPR50 in blue. Statistical comparisons by linear model with Šidák-corrected pairwise contrasts (emmeans); ***p < 0.001, **p < 0.01, *p < 0.05, ns = not significant. **(D)** Yeast complementation assay of WT NeRh50 in the ammonium-transport-deficient S. cerevisiae triple-mepΔ strain. Cells were plated onto minimal medium containing low ammonium as sole nitrogen source, with glutamate used as a positive control. Empty vector and E. coli AmtB were included as negative and positive controls, respectively.

To validate NeRh50 transport function *in vivo*, we expressed NeRh50 WT, *E. coli* AmtB WT, or pDR195 empty vector in the ammonium-transport-deficient *S. cerevisiae* strain 31019b (triple-mepΔ) and assessed growth on 0.5, 1, or 3 mM NH_4_^+^ as the sole nitrogen source. Glutamate was used as a non-selective nitrogen source control. All constructs grew robustly on glutamate, confirming viability and non-toxic expression (Figure 4D). At 0.5 mM NH_4_^+^, AmtB WT restored clear colony growth, whereas pDR195 produced no detectable colonies. NeRh50 WT failed to complement at day 3 but showed delayed colony formation by day 4. This behaviour is consistent with Michaelis–Menten estimates based on the SSME-derived apparent K_m_ for NeRh50 (∼59 mM, Figure 5B): at 0.5 mM NH_4_^+^, NeRh50 would be expected to operate at only ∼0.8% of its maximal transport capacity, compared with ∼38% for AmtB using the published K_m_ of ∼0.8 mM also determined using SSME^8^. At 1 mM NH_4_^+^, both AmtB WT and NeRh50 WT supported visible growth above the empty-vector background, indicating transporter-dependent ammonium acquisition. At 3 mM NH_4_^+^, all constructs produced colonies, including pDR195. This residual growth was unexpected as strain 31019b has previously been reported to fail to grow below 5 mM NH_4_^+^, albeit on a different medium^25,26^. Together, the yeast complementation data establish NeRh50 as a functional ammonium transporter *in vivo.* The delayed growth at 0.5 mM NH_4_^+^ and clear complementation at 1 mM NH_4_^+^ is suggestive of low-affinity transport behaviour.

**Figure 5.**
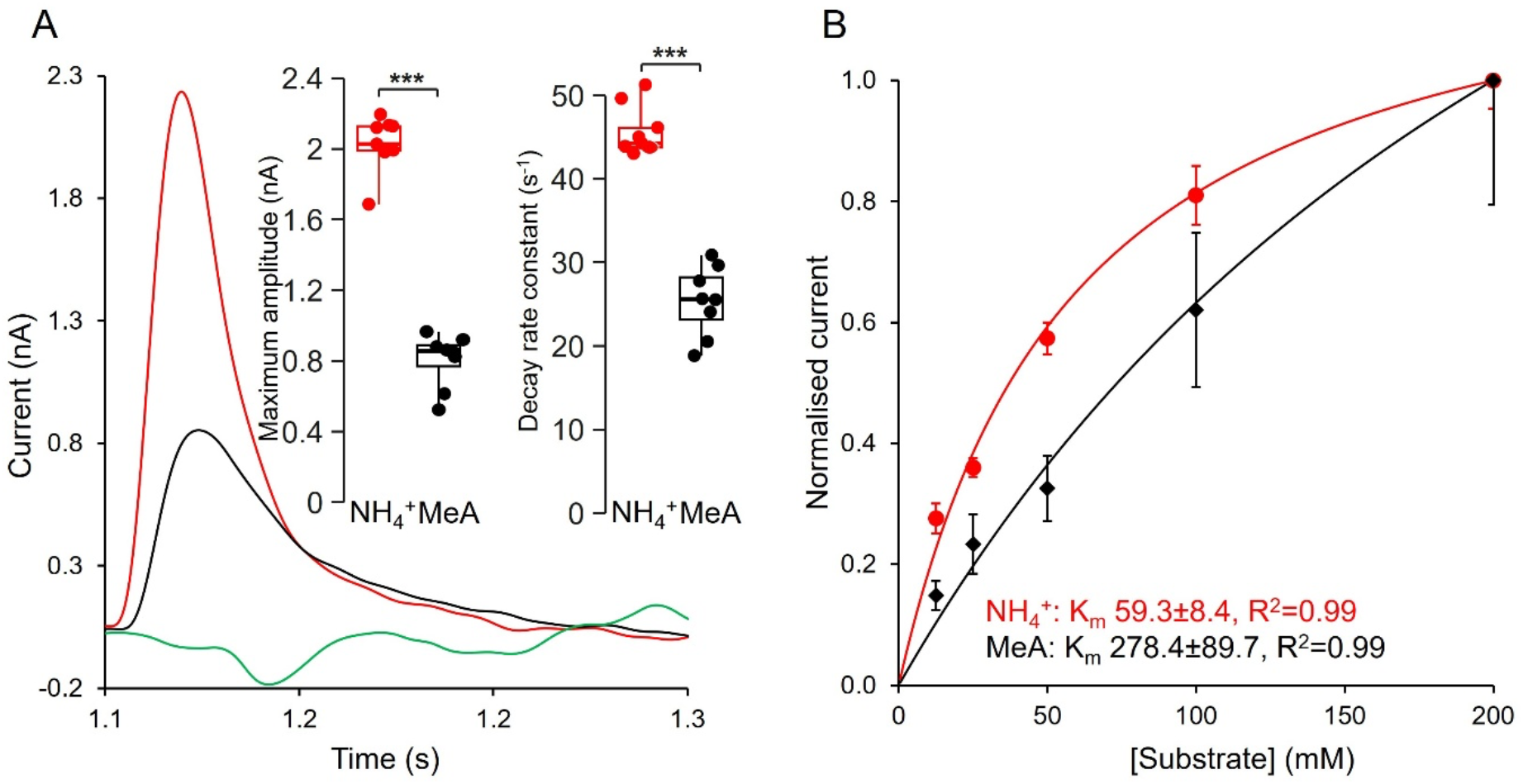
Substrate selectivity and kinetic analysis of wild-type NeRh50 by SSME. **(A)** Representative SSME transient currents recorded from proteoliposomes containing wild-type NeRh50 reconstituted at LPR5, following application of 200 mM NH₄⁺ (red), MeA (black), or K⁺ (green). Insets: maximum amplitude and decay rate constant k for NH₄⁺ and MeA. Individual data points are overlaid on box plots (median, interquartile range, range). Statistical comparisons by linear model with Šidák-corrected pairwise contrasts (emmeans); ***p < 0.001, **p < 0.01, *p < 0.05, ns = not significant. **(B)** Concentration–response curves for NH_4_^+^ and MeA transport by wild-type NeRh50, fitted to the Michaelis– Menten equation. Current amplitudes were normalised to 1.0 for comparison. Apparent kinetic constants are indicated on the panel. Data are presented as mean ± SD.

### NeRh50 is selective for ammonium and displays low-affinity saturable transport

The substrate selectivity of NeRh50 was assessed by replacing NH_4_^+^ with either K^+^ or methylammonium (MeA) at a fixed concentration of 200 mM (Figure 5A). A K^+^ pulse produced no detectable current, confirming that the transient current observed with NH_4_^+^ does not reflect a non-specific cation conductance. By contrast, MeA, which is widely used as a tracer of ammonium transport *in vivo*, elicited a clear signal, but with an amplitude of only ∼40% of that generated by NH_4_^+^ at the same concentration and with a markedly lower decay rate constant (amplitude and decay both Šidák-corrected p < 0.001; Figure 5A), indicating that it is transported substantially less efficiently. Concentration–response experiments performed at LPR 5 over the range 12.5–200 mM for both NH_4_^+^and MeA were fitted to the Michaelis–Menten equation (Figure 5B). Saturation was not reached over the concentration range tested, particularly for MeA. The fits yielded an apparent *K*m of 59 mM for NH₄⁺ and an estimated apparent *K*m of 278 mM for MeA, indicating an approximately five-fold lower apparent affinity for MeA. (Figure 5B). Thus, although MeA can be translocated by NeRh50, it is a relatively poor substrate analogue for the wild-type transporter. These data identify NeRh50 as a selective, low-affinity ammonium transporter and demonstrate that, as for Amt/Mep proteins, MeA is an imperfect surrogate for NH₄⁺ transport in the Rh family^6,13^. The low affinity of NeRh50 relative to AmtB, whose apparent K_m_ for NH₄⁺ is ∼0.8 mM^8^, is consistent with the physiology of *N. europaea*, an ammonia-oxidising chemolithoautotroph that can grow under millimolar ammonium conditions and whose niche is shaped by ammonium availability^27^.

### NeRh50 transport is pH dependent but independent of the proton gradient

To further define the energetic profile of NeRh50, its activity was examined by SSME over the pH range 5–8. Because pH shifts could, in principle, affect the stability of the transporter once reconstituted into proteoliposomes, a control experiment was first performed in which NeRh50 activity was measured at pH 7 before and after exposure to pH 5, 6, and 8 (Figure 6A-B). The transient current amplitude recorded at pH 7 remained comparable before and after exposure to the different pH conditions, and no appreciable change in the corresponding k values was observed. These data show that short exposure to acidic or basic pH does not irreversibly inactivate NeRh50 in proteoliposomes and therefore validate the use of this assay to examine pH-dependent effects on transport.

**Figure 6.**
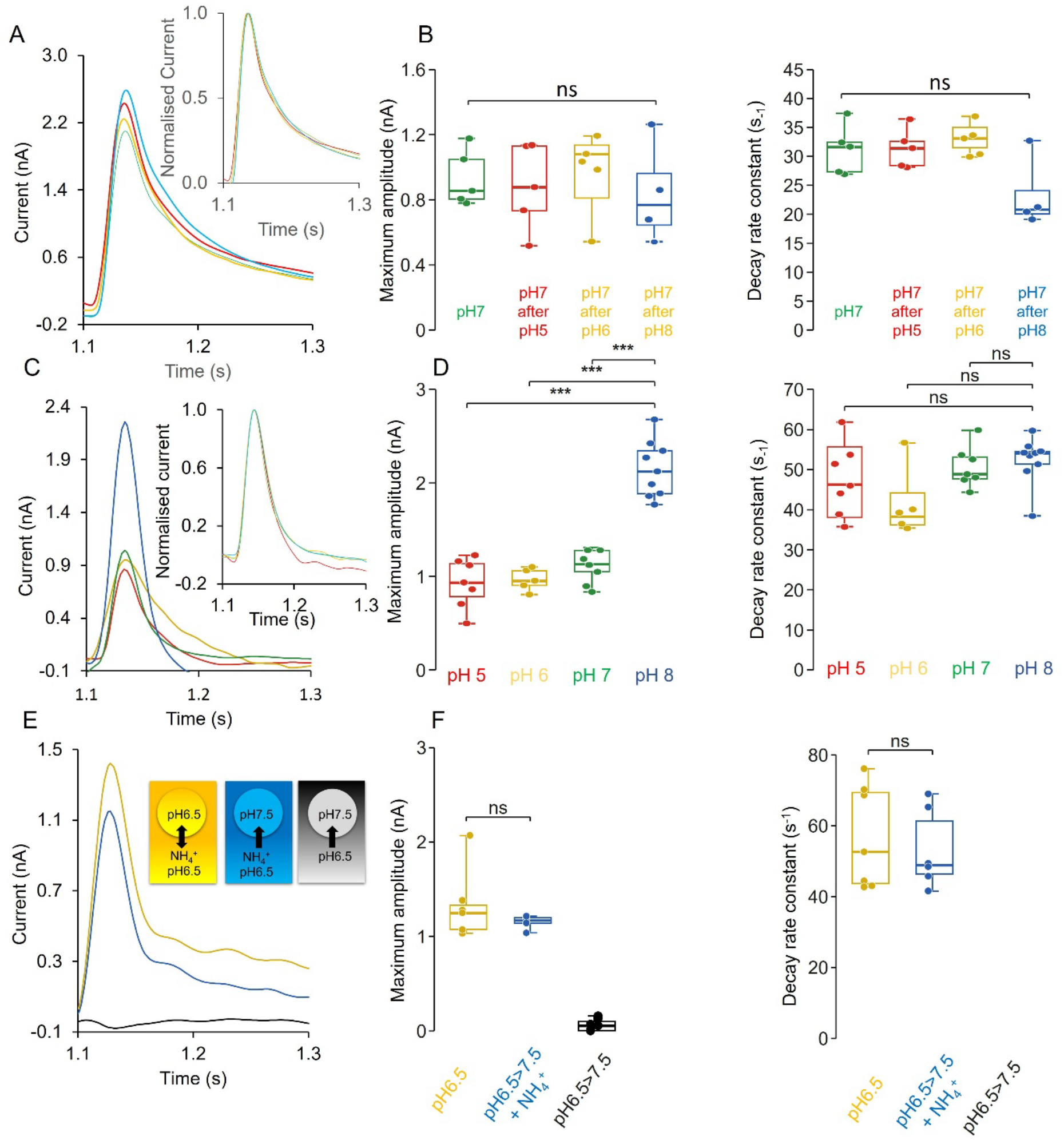
pH control, pH dependence, and proton-gradient experiment for wild-type NeRh50 measured by SSME. **(A)** Transient current measured after a 200 mM ammonium pulse in proteoliposomes containing NeRh50 at LPR10, at pH7 (green), pH7 after pH5 (red), pH7 after pH6 (yellow), and pH7 after pH8 (blue). Inset: peak-normalised traces for comparison. **(B)** Maximum amplitude and decay rate constant for the conditions in (A). Individual data points are overlaid on box plots (median, interquartile range, range). Statistical comparisons by linear model with Šidák-corrected pairwise contrasts (emmeans); ***p < 0.001, **p < 0.01, *p < 0.05, ns = not significant. **(C)** Representative SSME transient currents recorded at pH5 (red), pH6 (yellow), pH7 (green), and pH8 (blue). The corresponding normalised traces are shown in the inset, and the apparent decay constants (k) obtained under each pH condition are indicated on the panel. **(D)** Maximum amplitude and decay rate constant for the pH conditions in (C). Individual data points are overlaid on box plots (median, interquartile range, range). Statistical comparisons by linear model with Šidák-corrected pairwise contrasts (emmeans); ***p < 0.001, **p < 0.01, *p < 0.05, ns = not significant. **(E)** Proton-gradient experiment. Representative SSME transient currents recorded in the presence of 200 mM NH₄⁺ under static pH (pH6.5, yellow), in the presence of an inward proton gradient (pH6.5→7.5, blue), and in the absence of NH₄⁺ under the same proton-gradient configuration (black). **(F)** Maximum amplitude and decay rate constant for the conditions in (E). Individual data points are overlaid on box plots (median, interquartile range, range). No decay rate constant is shown for the pH6.5>7.5 (no NH₄⁺) condition, as these traces were flat with no measurable current from which to fit a decay. Statistical comparisons by linear model with Šidák-corrected pairwise contrasts (emmeans); ***p < 0.001, **p < 0.01, *p < 0.05, ns = not significant.

Having established this, transport activity was then measured directly at pH 5, 6, 7, and 8 at LPR 10 (Figure 6C-D). The current amplitude was similar at pH 5, 6 and 7 (0.93 ± 0.26 nA, 0.97 ± 0.12 nA and 1.12 ± 0.17 nA, respectively; all pairwise comparisons ns) but significantly higher at pH 8 (2.15 ± 0.30 nA; Šidák-corrected p < 0.0001 versus each of pH 5, 6 and 7). The apparent decay constant *k* did not differ significantly across the pH series (pH 5: 47.5 ± 12.3 s⁻¹; pH 6: 42.1 ± 9.9 s⁻¹; pH 7: 50.6 ± 5.1 s⁻¹; pH 8: 52.3 ± 5.9 s⁻¹; all pairwise comparisons ns). Thus, NeRh50 remains active throughout the pH 5–8 range, with alkaline pH selectively increasing the amplitude of the electrogenic current without a corresponding change in the kinetics of the transport cycle.

We next tested whether NeRh50 activity depends on the proton gradient itself, as expected for an NH_4_^+^/H⁺ symport mechanism. An inwardly directed proton gradient was imposed and compared with static pH 6.5 conditions in the presence of 200 mM NH_4_^+^ (Figure 6E-F). No appreciable difference in current amplitude or *k* was observed (both ns, p > 0.1; Figure 6F), arguing against stimulation by the bulk proton gradient and therefore against NH_4_^+^ /H⁺ symport. In a complementary control, the same inward proton gradient was applied in the absence of ammonium. No current was detected, excluding proton translocation through NeRh50 in the absence of substrate.

Together, these data show that NeRh50-mediated ammonium transport is pH-dependent but not driven by the proton gradient. This behaviour is consistent with the physiology of *N. europaea*, whose ammonia-oxidising activity is strongly influenced by external pH and favoured under neutral to alkaline conditions^28^. It also agrees with the thermodynamic framework proposed for AmtB, in which ammonium uptake must be active and membrane-potential-driven to accumulate intracellular NH_4_^+^ above the extracellular concentration^29^ and later discussed for the Amt/Mep/Rh superfamily^5^.

## Discussion

The Amt/Mep/Rh protein fold is highly conserved through evolution^4^, yet there are fundamental structural and functional differences between these transport proteins. Comparison of membrane environment and transport behaviour between Amt and Rh proteins using an integrated approach reveals significant divergence that underpins the physiological and ecological roles of these two protein family members.

The cryo-EM structure is consistent with the absence of the S1 periplasmic ammonium-binding site that contributes to substrate recruitment in AmtB, providing a structural context for the low apparent affinity measured by SSME. NeRh50 displayed an apparent *K*m of ∼59 mM for ammonium, approximately 70-fold higher than that reported for AmtB, while MeA was transported with an apparent *K*m of ∼278 mM ^30,8,31^. Imposing an inward proton gradient had no effect on transport, arguing against ΔpH-driven activity; the data are instead consistent with electrogenic, membrane-potential-dependent ammonium transport. This interpretation is consistent with the observation that ammonium accumulation in *N. europaea* cells is abolished by uncouplers^32^.

Cryo-EM also reveals lipid density at the monomer–monomer interface, with tight lipid retention supported independently by native MS and molecular dynamics simulations. This density occupies a position equivalent to the functionally important junction lipid site of AmtB^13,12^ but potentially adopts an inverted headgroup orientation reminiscent of lipid-transporting proteins^33^. Notably, this inverted orientation was not reproduced in our MD simulations, and its physiological relevance therefore remains uncertain. It may represent a lipid configuration captured during detergent extraction or, alternatively, a less favoured state that is not sampled spontaneously on the timescale of the simulations. Nevertheless, the convergence of cryo-EM, native MS and MD provides independent evidence for persistent lipid association at the NeRh50 monomer interface. Moreover, amphipathic molecules occupy equivalent interfacial regions in other members of the Amt/Mep/Rh superfamily, suggesting that this region represents an adaptable membrane-facing feature of the conserved trimeric architecture. We therefore propose that tightly associated lipids at the NeRh50 monomer interface may contribute to protein stability and/or function, while the precise functional significance and physiological orientation of these lipids remain to be established.

The transport properties of NeRh50 are also consistent with the physiological niche of *N. europaea*. This organism is associated with ammonium-replete environments and is less competitive at low ammonium than oligotrophic ammonia oxidisers^27^. The low apparent affinity of NeRh50 is therefore consistent with transport under substrate-rich conditions rather than high-affinity scavenging. The contrast with ammonia-oxidising archaea, whose affinities can reach the nanomolar range, illustrates how nitrogen acquisition strategies track ecological niche across orders of magnitude^34^. The pH dependence of NeRh50 transport further mirrors the pH-sensitive ammonia-oxidising activity of *N. europaea*, which is favoured under neutral to alkaline conditions^28^. Together with the absence of stimulation by an imposed proton gradient, these properties suggest that NeRh50 is adapted to support electrogenic ammonium transport under the substrate and pH conditions characteristic of *N. europaea* physiology.

The regulation and directionality of NeRh50 remain unresolved, specifically whether NeRh50 functions primarily as an importer, like canonical Amt proteins, or can also mediate ammonium export, as observed for human Rh proteins^35^. This question is particularly relevant because ammonium transport in many bacteria is controlled by an Amt–PII module, in which PII proteins regulate transporter activity to limit excessive influx, NH₄⁺/NH₃ cycling and ammonium toxicity^36^. *N. europaea* is unusual in this respect, as it lacks canonical PII-encoding genes and the PII uridylyltransferase GlnD^37^. One possibility is that the lower affinity of NeRh50, together with potential bidirectional Rh-type transport as observed for eukaryotic Rh proteins, provides an alternative strategy for ammonium homeostasis that reduces dependence on PII-mediated transporter gating. However, this hypothesis is unlikely to apply universally: *Nitrosospira multiformis* encodes Rh50 but lacks Amt while retaining GlnB and GlnD, whereas *Ca. Kuenenia stuttgartiensis* encodes Rh50 together with multiple Amt proteins and several GlnK/PII regulators^38,39^. A systematic comparative-genomics analysis will therefore be required to determine whether bacterial Rh50 acquisition, proposed to derive from a eukaryotic lineage^10^, is associated with loss, retention or rewiring of PII-based nitrogen sensing. Establishing whether NeRh50 can mediate ammonium efflux as well as import, and under which physiological conditions, will be important for defining how this unusual Rh-family transporter contributes to ammonium homeostasis in *N. europaea*.

Overall, our results show that NeRh50 combines a distinctive lipid-associated architecture with selective, low-affinity electrogenic ammonium transport. The convergence of cryo-EM, native mass spectrometry and molecular dynamics identifies the monomer interface as a site of persistent lipid association, while SSME and *in vivo* complementation establish NeRh50 as a functional ammonium transporter with transport properties markedly different from canonical AmtB. At the same time, the compatibility of NeRh50 transport with the charge-coupling framework proposed for AmtB raises the possibility that fundamental principles of electrogenic ammonium transport have been retained despite substantial divergence in substrate affinity, membrane-associated architecture and physiological context. Thus, diversification within the Amt/Mep/Rh superfamily may involve adaptation of a conserved structural and energetic framework to distinct membrane environments and physiological demands.

## Materials and Methods

### Protein expression and purification

NeRh50-(His)₆ was heterologously overexpressed in *E. coli* GT 1000 (Δ*glnK*, Δ*amtB*) using the pAD7 vector as previously described ^8^. Cells were grown in M9 minimal medium supplemented with 0.2 mg/mL L-glutamine as sole nitrogen source at 30°C, 210 rpm for 19 h. Cells were harvested via centrifugation at 7000 RCF for 20 min and resuspended in 50 mM Tris, 500 mM NaCl, 10% glycerol; pH 8 supplemented with 10 µg/mL DNAse and 200 µM PMSF. Membranes were isolated by French press lysis (20 kPSI, three passes) followed by differential centrifugation (27,000 RCF, 30 min, 4°C; 205,000 RCF, 60 min, 4°C) to remove cell debris and pellet inner membranes, respectively. Membranes were solubilised in 50 mM Tris-HCl pH 8.0, 500 mM NaCl, 10% glycerol, 2% lauryldimethylamine-N-oxide (LDAO) at 4°C for 2 h with gentle agitation. Insoluble material was removed by centrifugation at 205,000 RCF for 45 min at 4°C. NeRh50 was purified by immobilised metal affinity chromatography (IMAC) on a 1 mL HiTrap HP cobalt column (GE Healthcare) using a 40–500 mM imidazole gradient, followed by size exclusion chromatography (SEC) on a Superdex S200 10/300 GL column (GE Healthcare) equilibrated in 50 mM Tris-HCl pH 7.8, 100 mM NaCl, 0.09% LDAO. Protein purity was assessed at each step by SDS-PAGE and confirmed by anti-His Western blotting using a GenScript primary antibody and an ALEXA FLUOR 790 goat anti-mouse secondary antibody (ThermoFisher Scientific), detected by Li-COR Odyssey imaging.

### Cryo-EM sample preparation and data collection

For cryo-EM grid preparation, NeRh50 maintained in 0.09% LDAO was loaded onto a Superdex 200 10/300 Increase column (Cytiva) in 20 mM Tris, 100 mM NaCl, 0.03% n-Dodecyl-β-D-maltoside (DDM); pH 8 and immediately concentrated in a 100 kDa MWCO centrifugal concentrator (Sartorius). 3 µL of 2.5 mg/mL NeRh50 was applied to glow discharged Quantifoil R1.2/1.3, 300 mesh, Cu grids. The grids were blotted using a Vitrobot Mark IV (ThermoFisher) maintained at 100% humidity and 4°C, for 3 seconds at a blot force of 3 before being plunged into liquid ethane.

Preliminary grid screening to assess sample quality was performed on a Talos Arctica (FEI) operating at 200 kV equipped with a Falcon IV direct electron detector (Thermo Fisher Scientific). High resolution cryo-EM data were collected using a Titan Krios G3 (FEI) operating at 300 kV equipped with a K3 direct electron detector (Gatan) and BioQuantum imaging filter. A total of 17,996 movies were acquired automatically in counted super resolution mode using EPU software (Thermo Fisher Scientific) at 105,000x magnification, corresponding to a calibrated pixel size of 0.835 Å. Movies were collected with a target defocus range of −2.2 to −0.8 µm at a total dose of 39.4 e^-^/Å^2^ fractionated into 40 frames.

### Cryo-EM data processing

Processing was performed with the following software: RELION 5.0.0, 4.0.17 ^40^, cryoSPARC 4.5.3 ^41^, Scipion 3.3.1 ^42^ Xmipp3 ^43^ and CCP-EM 1.6.0 ^44^ with the overall workflow summarised in (Figure S4). All refinements and operations were performed in symmetry point C1 unless otherwise stated. RELION classification and refinement jobs were monitored using relion_tracker (https://github.com/bfcoop/relion_tracker) ^45^. Global resolution was estimated from gold-standard Fourier shell correlations (FSCs) using the 0.143 criterion, and local resolution estimation was calculated within RELION.

Raw movies were clustered into 68 optics groups using the extended metadata files of the EPU-data collection and the EPU_group_AFIS.py script (https://github.com/DustinMorado/EPU_group_AFIS). Motion correction and dose-weighting was performed using RELION’s implementation of MotionCor2 ^46^ and the contrast transfer function (CTF) subsequently estimated with CTFFIND4 ^47^. The default resnet8_u32 Topaz ^48^ model was used to pick a random subset of 500 micrographs yielding 146,468 particles at an FOM threshold of 0. Unwanted particles and contamination were removed from the particle stack through 2D classification in RELION at 3.34 Å/px. Four well defined classes were used to re-train the Topaz picking model which identified 6,600,550 particles (FOM threshold of −2) from the entire dataset. Two rounds of 2D classification were performed in RELION at 3.34 Å/px, yielding 2,121,388 particles which were imported into Cryosparc and subject to a further round of 2D classification.

1,833,579 particles from the best resolving classes were used for multi-class ab-initio reconstruction (K=3). 956,469 particles from the most populated class were re-extracted at 0.835 Å/px and subjected to non-uniform refinement ^49^ against their corresponding volume generating a 3.38 Å map. The particle stack was further cleaned via three rounds of heterogeneous refinement utilising the refined map and three noise-derived decoy volumes. Non-uniform refinement of the resulting particles (535, 893) yielded a 3.3 Å map before being imported into RELION using the cryosparc2star.py script from UCSF pyem ^50^.

3D classification with BLUSH regularisation ^51^ (K=6) yielded a single class with enhanced side chain density. The corresponding 177,850 particles were subsequently subjected to CTF refinement ^52^ and Bayesian polishing ^53^ in an expanded box (400 px) before 3D auto-refinement in RELION which yielded a 3.1 Å map. Two further identical RELION 3D auto-refine runs were performed in parallel, following the strategy proposed by ^54^ to reduce bias and variance. Particle alignments from all three runs were compared pairwise using Xmipp compare angles and particles showing angular and shift differences greater than 1° and 1 pixel, respectively were discarded.

The resulting stack, of 164,696 particles, was subjected to a final RELION 3D-auto refinement and sharpened using RELION post-processing, resulting in the final 3.06 Å NeRh50 map (EMD-59467). The 3D-auto refinement and post-processing steps were repeated with a soft mask, excluding density corresponding to the C-terminal helical moieties, yielding the final 3.03 Å NeRh50^CORE^ map (EMD-59468). Repeating these steps with the application of C3 symmetry yielded the final 2.88 Å NeRh50^CORE-C3^ map (EMD-59469).

### Atomic model building and refinement

Modelangelo ^55^ (implemented in RELION 5.0) was used to build initial protein models into RELION post-processed maps. The resulting models were refined in real space using Phenix ^56,57^ interspersed with manual inspection and adjustment in Coot ^58^. All models were validated using MolProbity ^59^ within Phenix.

### Structural analysis and figure preparation

Investigation and comparison of cryo-EM and X-ray derived models was carried out using UCSF ChimeraX ^60^. Crystallographic coordinates for NeRh50 (PDBs: 3B9W, 3B9Y and 3B9Z), *Ec*AmtB (PDBs: 4NH2 and 6B21), *Ca*Mep2 (PDBs: 5AEZ and 5AH3) and *Hs*RhCG (PDB 3HD6) were downloaded from the PDB. To facilitate interpretation of cryo-EM density within figures, selected maps were filtered using LAFTER ^16^. Maps processed in this manner are clearly indicated in the corresponding figure legends. The masks used during LAFTER filtering were the same as those employed for post-processing in RELION. Figures were prepared using UCSF ChimeraX and Inkscape.

### Native mass spectrometry

Purified NeRh50 at 32 µM was buffer-exchanged into “MS buffer” containing 200 mM ammonium acetate (pH 8.0) supplemented with 1.0% n-Octyl β-D-glucopyranoside (β-OG) using a Micro Bio-Spin 6 desalting column (Bio-Rad). The desalted protein was further diluted to a final concentration of 5.0 µM and about 3 μL aliquot was transferred into a gold-coated borosilicate capillary prepared in-house and was mounted on the nano ESI source of a Q-Exactive hybrid quadrupole-Orbitrap mass spectrometer (Thermo Fisher Scientific, Bremen, Germany). The instrument settings were 1.2 kV capillary voltage, S-lens RF 200%, argon UHV pressure 3.3 × 10_−10_ mbar, the capillary temperature was set to 200 °C, and resolution of the instrument was set to 17,500 at a transient time of 64 ms. The noise level was set at 3 rather than the default 4.6. Voltages of the ion transfer optics –injection flatapole, inter-flatapole lens, bent flatapole, and transfer multipole were set to 5, 3, 2, and 30 V respectively. Ions were activated in the HCD cell at 100 V without in-source trapping.

### Molecular dynamics simulations

All molecular dynamics simulations were carried out using GROMACS 2021.5 ^61^ with the CHARMM36m force field ^62^ supplemented by the CHARMM-WYF extension ^63,64^ for treatment of cation-π interactions. To improve computational efficiency, hydrogen mass repartitioning (HMR) ^65,66^ was applied to the protein and lipid bilayer during the production simulations.

The transmembrane region of NeRh50 was embedded in a lipid bilayer designed to mimic the composition of the *E. coli* inner membrane. Both membrane leaflets consisted of 75% 16:0/18:1 palmitoyloleoyl-phosphatidylethanolamine (POPE), 20% 16:0/18:1 palmitoyloleoyl-phosphatidylglycerol (POPG), and 5% cardiolipin. The NeRh50–membrane system was constructed using the CHARMM-GUI Bilayer Builder ^67,68^. One NeRh50 was incorporated into a bilayer with lateral dimensions of 15 × 15 nm², with a 5 nm-thick aqueous layer placed on each side of the membrane. The resulting system was neutralized and adjusted to a NaCl concentration of 0.15 M.

Prior to equilibration, the system was energy-minimized using the steepest-descent algorithm until the maximum force decreased below 1000 kJ mol⁻¹ nm⁻¹. The minimized system was subsequently equilibrated through six consecutive stages. The first three equilibration stages were each run for 1.25 ns with a timestep of 1 fs, whereas the following three stages were each performed for 5 ns using a 2 fs timestep. Position restraints were imposed on the protein and membrane during equilibration, with the corresponding force constants progressively reduced across successive stages. The temperature was maintained at 313 K using the velocity-rescaling thermostat ^69^, and the pressure was kept at 1 bar using the C-rescale barostat ^70^. Long-range electrostatic interactions were treated using the Particle Mesh Ewald (PME) method ^71^, while a cutoff distance of 1.2 nm was used for short-range electrostatic and van der Waals interactions. Bonds involving hydrogen atoms were constrained with the LINCS algorithm ^72^.

Following equilibration, production simulations were conducted for 1 μs using a 4 fs timestep. Three independent replicates were performed, with each replicate independently subjected to the complete energy-minimization and equilibration procedure.

### Simulations analysis

Molecular graphics and structural representations were generated using ChimeraX v1.11.1 _60_. Lipid–protein interactions were analysed from the molecular dynamics trajectories using PyLipID ^17^, with particular emphasis on lipid residence time as a measure of the kinetic stability of lipid–protein interactions. For each lipid species, all trajectory frames and all lipid atoms were included in the analysis, with lipid–residue contacts defined using a dual-distance cutoff of 0.475 and 0.70 nm, where the shorter cutoff defines contact formation and the longer cutoff maintains an existing interaction, thereby reducing artificial fragmentation of binding events caused by fluctuations around a single distance threshold. Longer residence times indicate more persistent lipid–protein interactions, whereas shorter residence times reflect more rapidly exchanging lipid contacts.

### Proteoliposome preparation

*E. coli* polar lipids and 1-palmitoyl-2-oleoyl-sn-glycero-3-phosphocholine (POPC; Avanti Polar Lipids) were mixed at 2:1 (w/w), dried under a stream of nitrogen and desiccated under vacuum for 2 hours. Lipids were rehydrated in liposome buffer (100 mM potassium phosphate pH 7.6, 300 mM KCl) at 5 mg/mL and extruded eleven times through a 100 nm polycarbonate membrane (Mini-Extruder, Avanti Polar Lipids) to produce unilamellar liposomes of defined size. The detergent saturation (R∼sat∼) and solubilisation (R∼sol∼) concentrations were determined empirically by successive addition of 1 µL aliquots of 25% Triton X-100 to 500 µL liposomes (5 mg/mL) with absorbance monitoring at 540 nm. Protein was incorporated at R∼sat∼ at lipid-to-protein ratios (LPR) of 5:1, 10:1 or 50:1 (w/w) by incubation at 25°C for 5 min then 4°C for 30 min. Detergent was removed by two consecutive incubations with Bio-Beads SM-2 (Bio-Rad; ∼0.2 g per 500 µL) at 4°C (2 h then overnight).

Proteoliposomes were washed three times by ultracentrifugation (200,000 × g, 60 min, 4°C), resuspended in liposome buffer at 5 mg lipid/mL, aliquoted and stored at −80°C. Successful protein insertion was confirmed by SDS-PAGE analysis of wash fractions and the final proteoliposome pellet.

### Solid-supported membrane electrophysiology

SSME measurements were performed using a SURFE²R N1 instrument (Nanion Technologies) as described ^8^. Briefly, 3 µL of proteoliposomes were applied to pre-hydrated SSME sensors, adsorbed by centrifugation at 2,000 × g for 30 min at 4°C and dried at room temperature for 30 min. Substrate-induced transient currents were elicited by rapid solution exchange between non-activating and activating solutions of matched ionic strength. Maximum current amplitudes and decay rate constants (k values) were extracted by mono-exponential fitting of the transient current decay. For LPR-dependence experiments, a 200 mM NH₄Cl pulse was applied at LPR 5, 10 and 50 to confirm that the transient current represents a complete translocation cycle rather than a substrate-binding event. For substrate selectivity measurements, activating solutions contained 200 mM NH₄Cl, methylammonium chloride (MeA) or KCl. Kinetic parameters were determined from concentration-response experiments (12.5–200 mM NH₄⁺ or MeA; LPR 5) fitted to the Michaelis-Menten equation; maximum amplitudes were normalised to 1.0 for comparison between substrates. For pH dependence experiments, measurements were carried out at pH 5, 6, 7 and 8 using matched solution pairs; protein stability was confirmed by recovery of pH 7 amplitude before and after exposure to each pH condition. For proton gradient experiments, proteoliposomes were prepared with an intra-liposomal pH of 7.5 and measurements were performed at an extra-liposomal pH of 6.5 in the presence or absence of 200 mM NH₄⁺; control measurements were performed at static pH 6.5 under identical conditions. Protein-free liposomes were used as negative controls throughout.

### Statistical analysis

All statistical comparisons were performed in R ^73^ using ordinary least-squares linear models (Value ∼ Group) fitted with the base R lm() function. For each comparison, model diagnostics (residual plots, assessed using the R performance package^74^) were used to check for heteroscedasticity, and HC3 heteroskedasticity-consistent standard errors ^75^ were applied throughout as a conservative default. Pairwise contrasts between group levels were estimated using the emmeans package ^76^ and corrected for multiple comparisons using the Šidák method ^77^. For experiments spanning multiple recording days (Figure 6F), a Day term was included in the model to account for batch variation. Box plots throughout report the median, interquartile range and full range of each group; individual data points are jittered for visibility. Significance is denoted as *** p < 0.001, ** p < 0.01, * p < 0.05, ns = not significant, matching the convention used in each figure legend.

### Yeast complementation assays

NeRh50 and *E. coli* AmtB were expressed from the pDR195 plasmid under the control of the PMA1 promoter and transformed into *S. cerevisiae* strain 31019b (MATa ura3 *mep1*Δ *mep2*Δ::LEU2 *mep3*Δ::KanMX2). Transformed cells were selected on YNB minimal medium lacking uracil and grown in YNB minimal buffered medium (pH 6.1, 3% glucose) supplemented with 1–3 mM (NH_4_)_2_SO_4_ or Glu (0.2%) as sole nitrogen source. Cultures were adjusted to OD 0.6 in HEPES buffer pH 6.1, washed three times, diluted 1:100 and 50 µL spread onto selective medium. Plates were imaged after 3 and 4 days of incubation at 30°C. Images were acquired as greyscale JPEG files and a linear contrast stretch was applied identically to all images by subtracting the 20th intensity percentile as the agar baseline and rescaling so that the 98th percentile mapped to maximum intensity, rendering the agar surface dark and colonies white.

## Data Availability

Cryo-EM reconstructions and corresponding coordinates have been deposited in the Electron Microscopy Data Bank and the PDB, respectively: NeRh50 (EMD-59467 and PDB 33QD), NeRh50^CORE^ (EMD-59468 and PDB 33QE) and NeRh50^CORE-C^^3^ (EMD-59469 and PDB 33QF). The raw cryo-EM dataset used in this study has been deposited to the Electron Microscopy Public Image Archive (EMPIAR) under accession code EMPIAR-13937.

---

**Supplementary Table 1:**
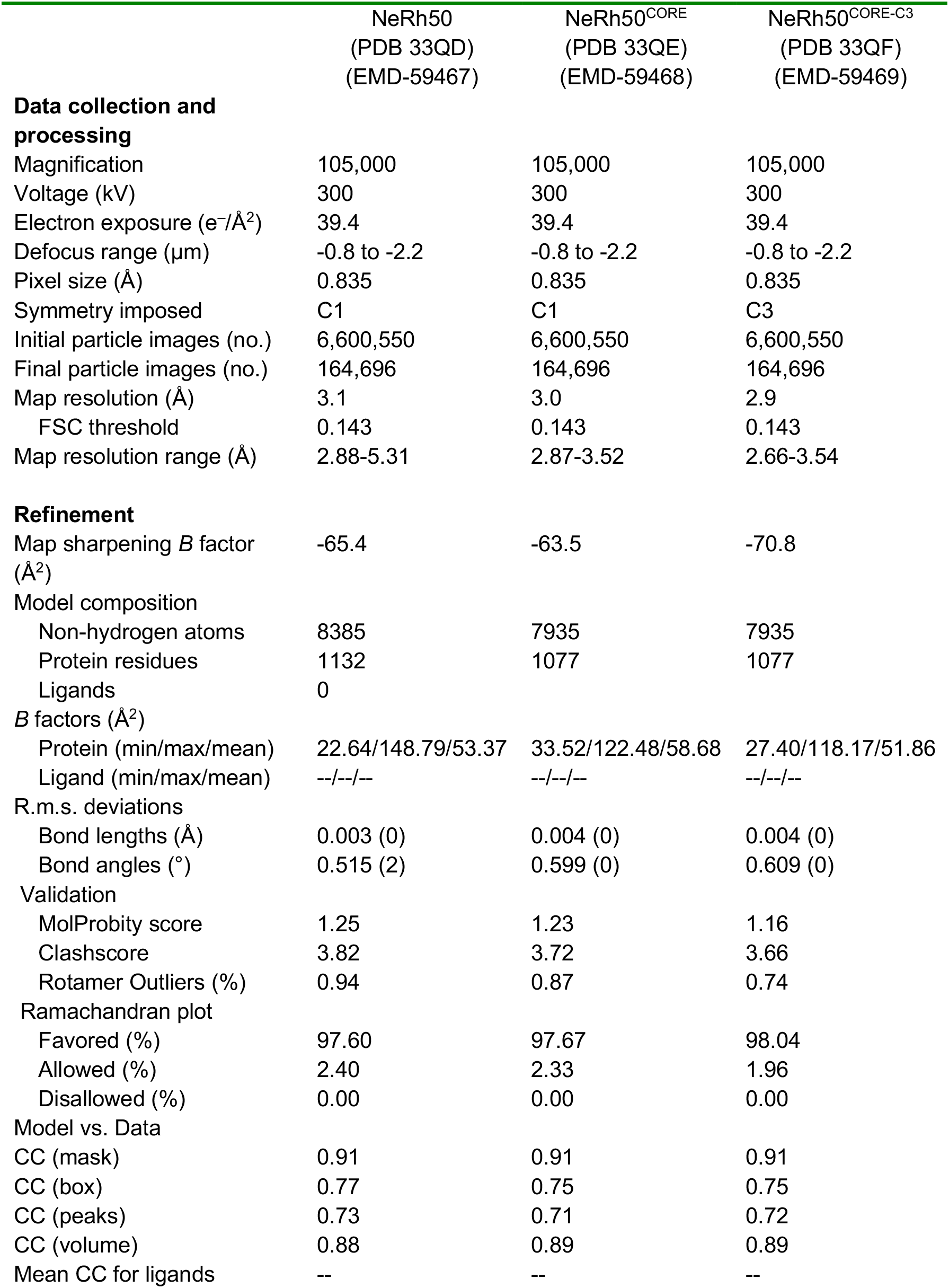
Cryo-EM data collection, refinement and validation statistics.

## Supplementary Figures

**Figure S1.**
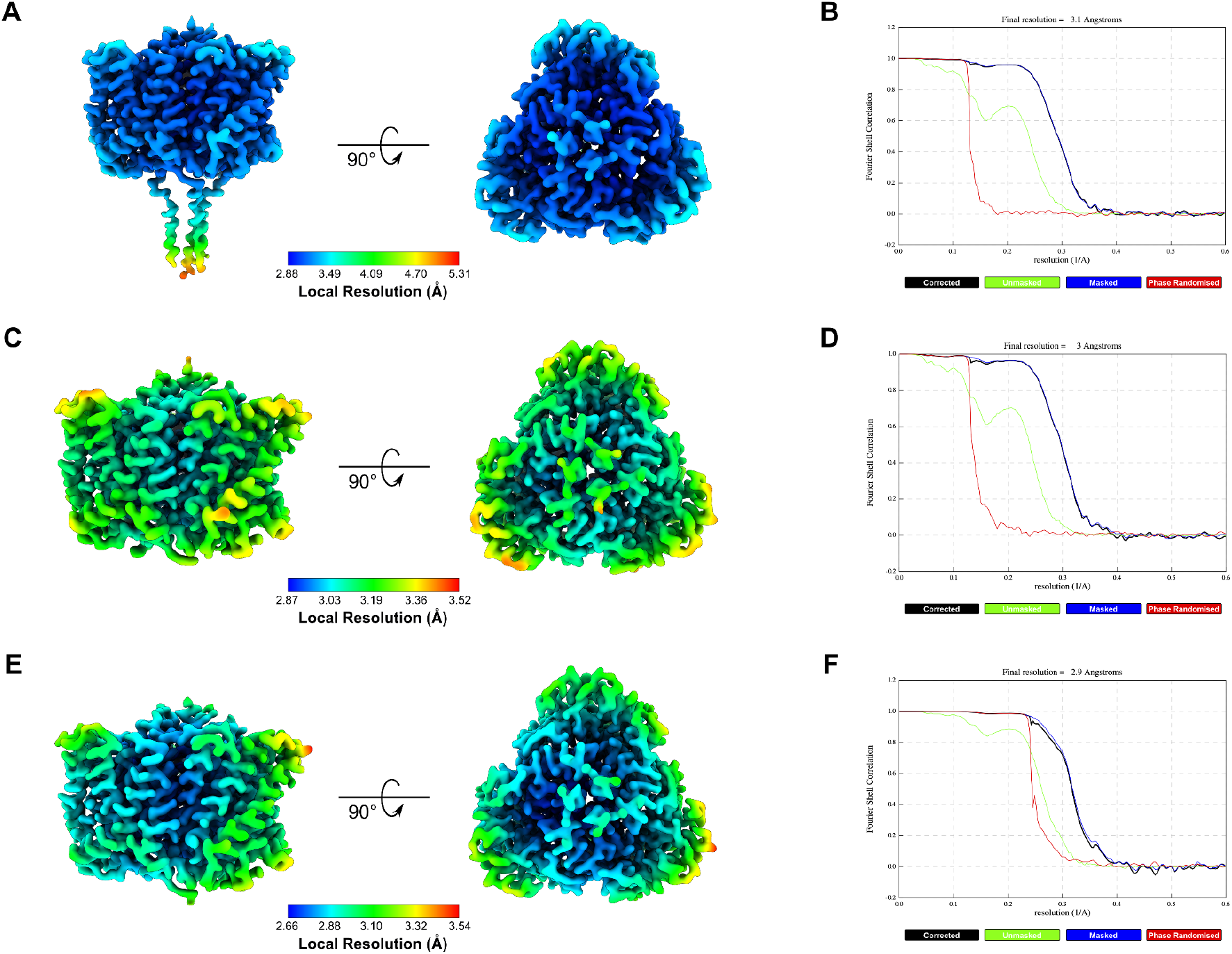
Resolution Validation of NeRh50 cryo-EM Maps. **(A)** LAFTER^16^ filtered reconstruction of NeRh50 at 3.1 Å coloured by local resolution (Å) with the corresponding colour key shown below the map. **(B)** Fourier shell correlation (FSC) curve for the NeRh50 cryo-EM reconstruction at 3.1 Å with global resolutions determined using the gold-standard 0.143 criterion. **(C)** LAFTER^16^ filtered reconstruction of the NeRh50 core at 3 Å coloured by local resolution (Å) with the corresponding colour key shown below the map. **(D)** Fourier shell correlation (FSC) curve for the NeRh50 core cryo-EM reconstruction at 3 Å with global resolutions determined using the gold-standard 0.143 criterion. **(E)** LAFTER^16^ filtered C3 reconstruction of the NeRh50 core at 2.9 Å coloured by local resolution (Å) with the corresponding colour key shown below the map. **(F)** Fourier shell correlation (FSC) curve for the C3 NeRh50 core cryo-EM reconstruction at 2.9 Å with global resolutions determined using the gold-standard 0.143 criterion.

**Figure S2.**
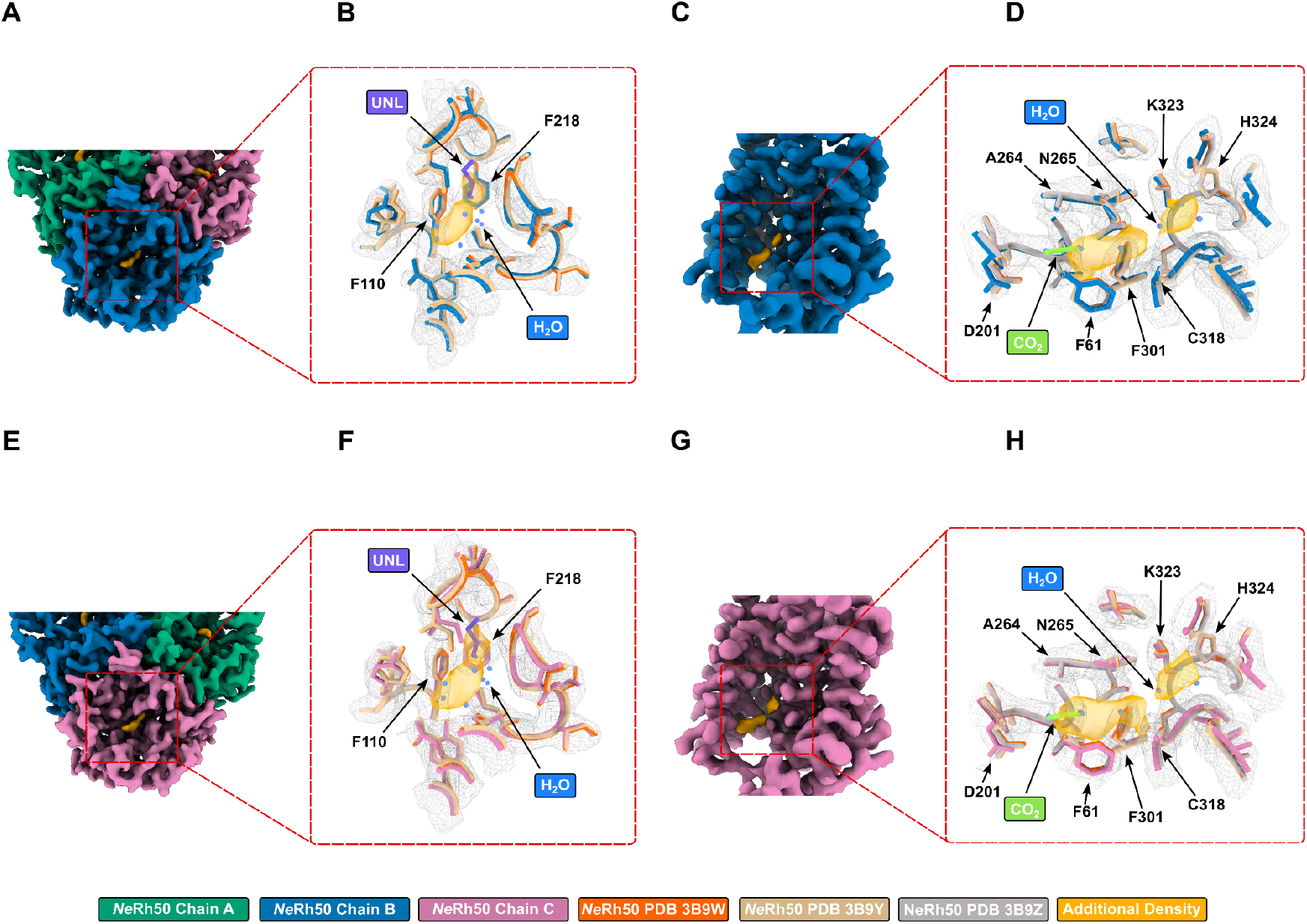
Additional Densities within the NeRh50 vestibules. **(A)** Elongated density (gold) present within the periplasmic vestibule of NeRh50. The Relion postprocessed map is contoured at 1.5 σ (map level 0.008), with density corresponding to chains A, B and C coloured green, blue and pink respectively. **(B)** Density (gold) present within the periplasmic vestibule of NeRh50. The Relion postprocessed map contoured at 1.5 σ (map level 0.008) and displayed as a grey mesh. NeRh50 models derived from our cryo-EM data, PDB 3B9W and PDB 3B9Y are coloured dark blue, orange and tan, respectively. Water molecules modelled in both PDBs 3B9W and 3B9Y are coloured blue, whilst the unknown ligand modelled in PDB 3B9Y is coloured purple. **(C)** Elongated density (gold) present within the cytoplasmic vestibule of NeRh50. The Relion postprocessed map is contoured at 1.5 σ (map level 0.008), with density corresponding to chain B coloured dark blue. Density corresponding to residues 32-82 (TM1-2) of NeRh50 is removed to aid clarity. **(D)** Densities (gold) present within the cytoplasmic vestibule of NeRh50. The Relion postprocessed map is contoured at 1.5 σ (map level 0.008) and displayed as a grey mesh. NeRh50 models derived from our cryo-EM data, PDB 3B9W, PDB 3B9Y and PDB 3B9Z are coloured dark blue, orange, tan and grey, respectively. Water molecules modelled in both PDBs 3B9W and 3B9Y are coloured blue, whilst the CO_2_ molecule modelled in PDB 3B9Z is coloured lime green. **(E)** Elongated density (gold) present within the periplasmic vestibule of NeRh50. The Relion postprocessed map is contoured at 1.5 σ (map level 0.008), with density corresponding to chains A, B and C coloured green, blue and pink respectively. **(F)** Density (gold) present within the periplasmic vestibule of NeRh50. The Relion postprocessed map contoured at 1.5 σ (map level 0.008) and displayed as a grey mesh. NeRh50 models derived from our cryo-EM data, PDB 3B9W and PDB 3B9Y are coloured pink, orange and tan, respectively. Water molecules modelled in both PDBs 3B9W and 3B9Y are coloured blue, whilst the unknown ligand modelled in PDB 3B9Y is coloured purple. **(G)** Elongated density (gold) present within the cytoplasmic vestibule of NeRh50. The Relion postprocessed map is contoured at 1.5 σ (map level 0.008), with density corresponding to chain B coloured pink. Density corresponding to residues 32-82 (TM1-2) of NeRh50 is removed to aid clarity. **(H)** Densities (gold) present within the cytoplasmic vestibule of NeRh50. The Relion postprocessed map is contoured at 1.5 σ (map level 0.008) and displayed as a grey mesh. NeRh50 models derived from our cryo-EM data, PDB 3B9W, PDB 3B9Y and PDB 3B9Z are coloured pink, orange, tan and grey, respectively. Water molecules modelled in both PDBs 3B9W and 3B9Y are coloured blue, whilst the CO_2_ molecule modelled in PDB 3B9Z is coloured lime green.

**Figure S3.**
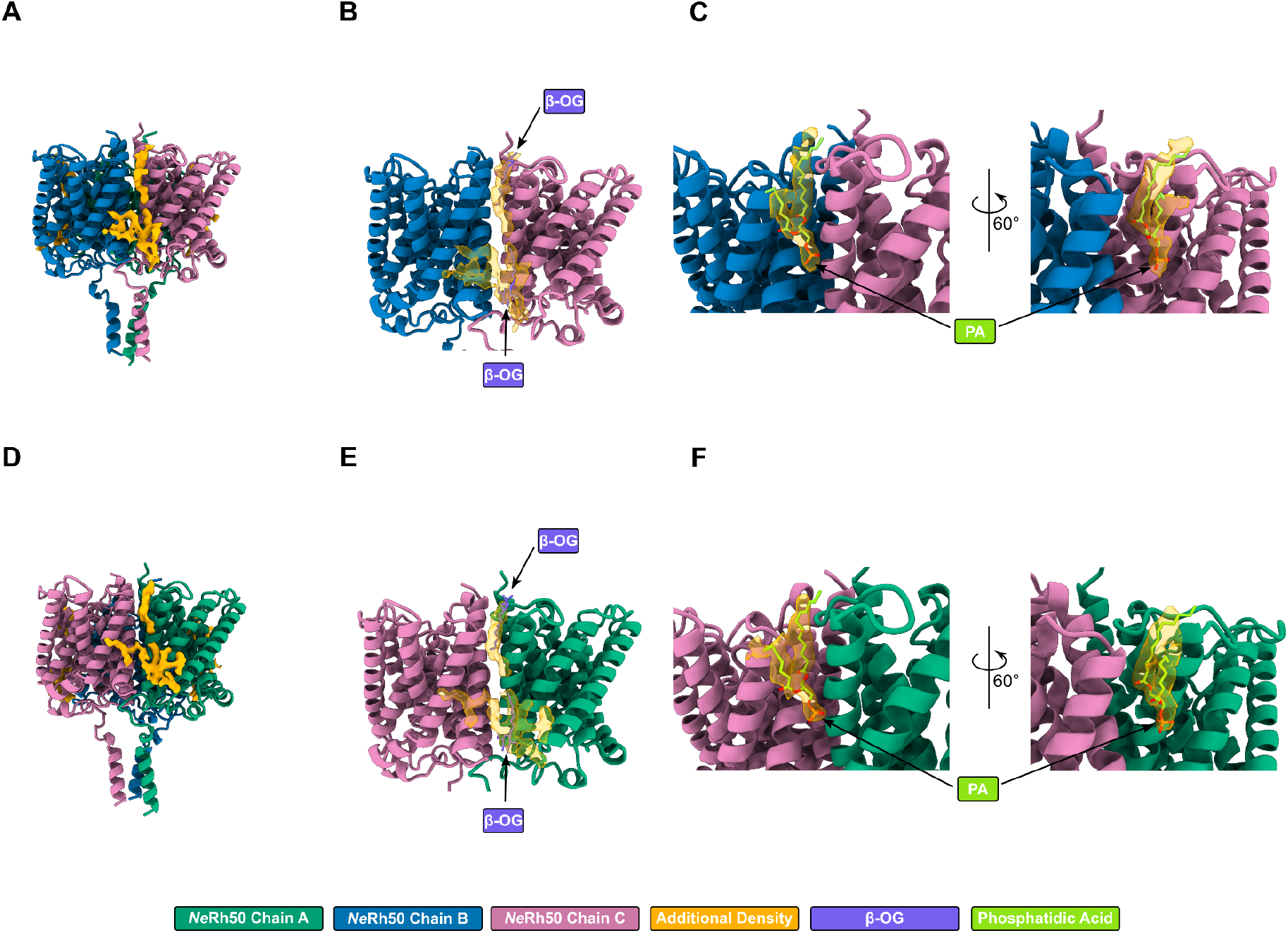
Annular densities surround the NeRh50 transmembrane region. **(A)** Annular densities (gold) located at the NeRh50 monomer interface between chains B and C (blue and pink cartoons respectively). **(B)** Interface density (transparent gold) between NeRh50 chains B and C (blue and pink cartoons, respectively) overlaid with the two β-OG molecules (purple) modelled in the crystallographic structure of NeRh50 (PDB 3B9Y). **(C)** Phosphatidic acid (PA, lime green) modelled into the periplasmic leaflet interface density (transparent gold) between NeRh50 chains B and C (blue and pink cartoons, respectively). **(D)** Annular densities (gold) located at the NeRh50 monomer interface between chains C and A (pink and green cartoons respectively). **(E)** Interface density (transparent gold) between NeRh50 chains C and A (pink and green cartoons, respectively) overlaid with the two β-OG molecules (purple) modelled in the crystallographic structure of NeRh50 (PDB 3B9Y). **(F)** Phosphatidic acid (PA, lime green) modelled into the periplasmic leaflet interface density (transparent gold) between NeRh50 chains C and A (pink and green cartoons, respectively).

**Figure S4.**
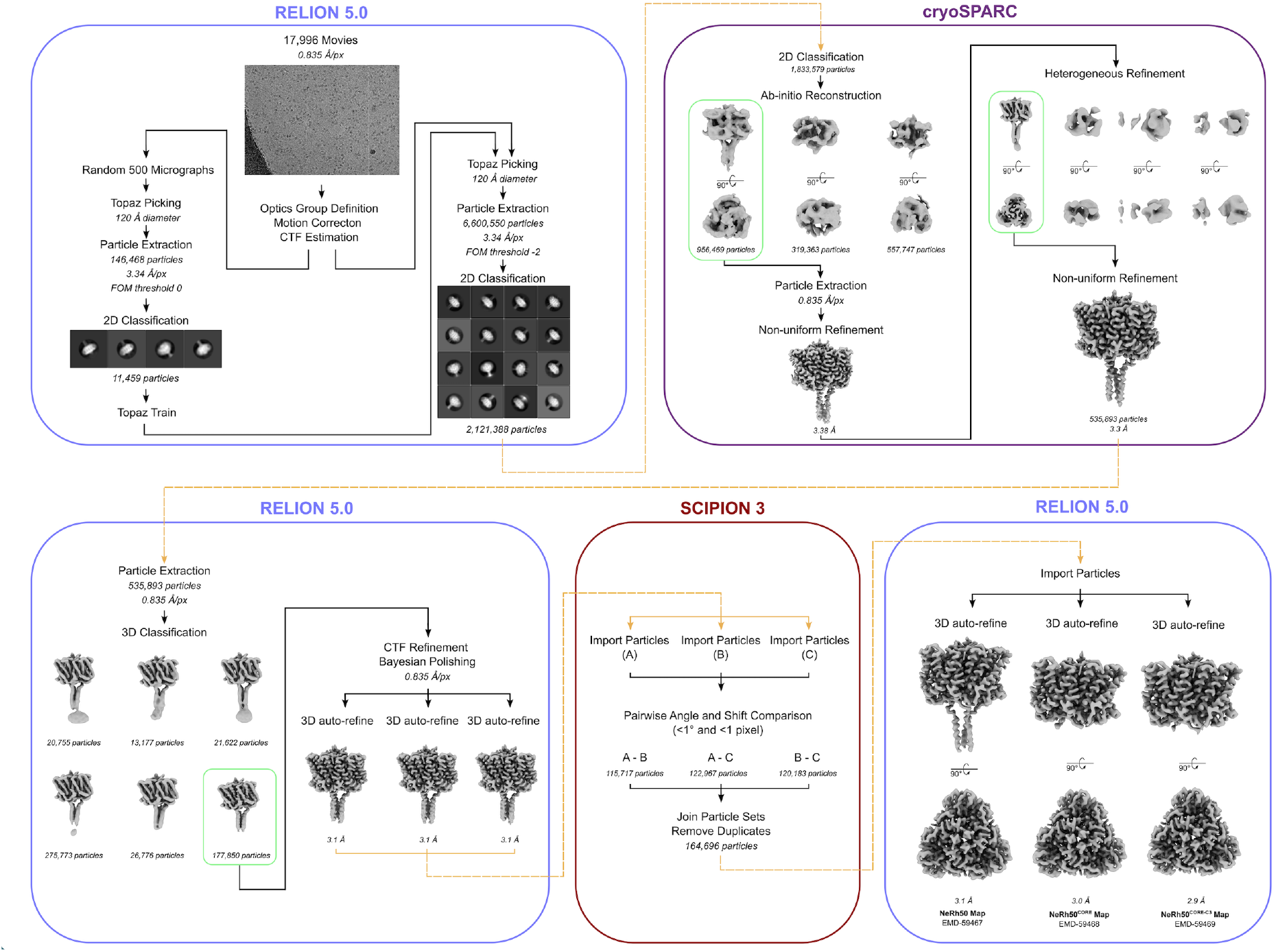
Cryo-EM processing of NeRh50. Cryo-EM data processing workflow for the NeRh50 sample in DDM.

